# Cell junction disruption drives translocation of gasdermin A and gasdermin B from cytoskeleton to plasma membrane during acantholysis

**DOI:** 10.64898/2026.08.17.745251

**Authors:** Kidong Kang, Ying Wang, Edward A Miao

## Abstract

Gasdermins (GSDMs) are a family of pore forming protein that trigger pyroptosis by permeabilizing cell membranes. Pyroptotic cells often release the proinflammatory cytokines interleukin-1β (IL-1β), and IL-18, thereby promoting an inflammatory response. GSDMs are typically cleaved by caspases or granzymes, which enable their translocation to the membrane. Here, we showed GSDMA and GSMDB localize to the cytoskeletal fraction of keratinocytes. Disruption of cell junctions causes gasdermin A and B (GSDMA and GSDMB) to translocate to the membrane fraction in the absence of cleavage. Cell junction disrupted keratinocytes release post-translationally modified keratins, but not IL-1β or IL-18. These events depend on endocytic mechanisms associated with recycling of cell junctional proteins. Our study suggests that cell junction disruption can drive translocation of GSDMA and GSDMB from cytoskeleton to plasma membrane in keratinocytes, however there may be a subsequent trigger that causes the conformational change allowing these gasdermins to form open pores.

## Introduction

Gasdermins (GSDMs) are a family of pore-forming proteins that play a pivotal role in innate immune defense against intracellular pathogens. GSDMs consist of N-terminal domain and a C-terminal domain connected by a linker region. Upon activation, caspases or granzymes cleave the proteolytic cleave linker, releasing the N-terminal domain to dock onto the membrane and form large oligomeric pores. For example, caspase-1 or −11 cleave the GSDMD linker to cause pyroptosis. In contrast, GSDME pore formation can occur without linker cleavage.

Ultraviolet-C irradiation has been reported to cause activation of GSDME, promoting its translocation to the membrane and pore formation, thereby triggering pyroptosis even in absence of a proteolytic cleavage event (1). Caspase-1 also cleaves the proinflammatory cytokines interleukin-1β (IL-1β) and IL-18 to their active forms, and these are released through the simultaneously activated GSDMD pores. GSDMs are named because some family members show specific expression patterns to the gastrointestinal tract (gas-) or to the epidermis of the skin (-dermin). For example, GSDMA is highly expressed in keratinocytes in the skin.

The skin forms a barrier between the body and the environment, with the most important barrier cells being the stratified and cornified epithelial cells called keratinocytes. Keratinocytes develop from a single basal layer into a stratified epithelium that forms a physical and immune barrier against injury, pathogen, and environmental stress. This barrier formation occurs as keratinocytes migrate from the proliferative basal layer, differentiate through the spinous layer, and reach terminal differentiation in granular layer; this process is known as cornification.

Keratinocytes depend on specialized cell junction complexes to maintain the mechanical integrity of the skin and facilitate signal transduction to coordinate cytoskeletal crosstalk. Crosstalk between cell adhesion complexes enables cells to sense and respond to physical forces through the regulation of cytoskeleton tension, thereby coordinating cell movement and development (2, 3). These adhesion complexes are categorized into cell-cell adhesion and cell-matrix adhesion. Cell-cell adhesion links adjacent keratinocytes together and plays a critical role in maintaining epithelial tissue integrity. Adhesion is primarily maintained by three major junctional structures: desmosomes, adherens junctions (AJs), and tight junctions (TJs).

Disruption of cell-cell adhesion is associated with numerous pathological conditions, for example autoimmune antibodies against desmoglein 3 causes a skin blistering disease called pemphigus vulgaris. Cell-matrix adhesion complexes anchor keratinocytes in the basal layer to the underlying extracellular matrix (ECM), called the basement membrane, to keep stratified keratinocytes of the epidermis tightly adhered to the dermis, which is critical for barrier integrity. Cell-matrix adhesion complexes are composed of hemidesmosome and focal adhesions. The homeostasis, integrity, and stability of these cell junctions are controlled through complex interactions with cytoskeletal proteins. Defects in these cell-matrix adhesion, for example caused by autoinhibits against the hemidesmosome protein BP130, causes another severe blistering diseases called bullous pemphigoid. Phosphorylation and dephosphorylation processes derived from specific protein kinase and phosphatases play an essential role in forming and maintaining the integrity of the multiprotein complexes that form desmosomes and hemidesmosomes. Protein Ser/Thr phosphatase 2A (PP2A) regulates epithelial cell-cell junctions, either directly dephosphorylating component proteins or indirectly regulating signaling pathway that contribute to junctional integrity. Dysregulation of PP2A activity consequently disrupts cell junctions and causes cytoskeletal dynamics (4, 5).

As they mature and move up in the stratified epithelial layers, keratinocytes undergo an obligatory programmed cell death that results in cornification. As keratinocytes move from the basal layer to higher epithelial layers, they express distinctive genes including GSDMA and GSDMB. These two genes are encoded adjacent to each other. GSDMA senses virulence factors from skin pathogens such as Group A *Streptococcus* and *Staphylococcus aureus* (6, 7).

Mice are unusual among mammals in that they have deleted GSDMB, and have triplicated GSDMA into *Gsdma1, Gsdma2, and Gsdma3*. *Gsdma1-3* triple knockout mice have normal development and no visually apparent defects in skin development or health. After skin damage, these genes have been reported to be required for keratinocyte cornification during regeneration (8). GSDMA is highly expressed in terminal differentiated keratinocytes (8, 9). GSDMB is expressed in human terminally differentiation keratinocyte and is also induced in psoriasis lesions (9, 10). Notably, *Gsdma1/a3*-deficient mice are healthy and have visually normal skin, however they exhibit a defect in epithelia repair after damage (8). GSDMB regulates focal adhesion kinase (FAK) phosphorylation to promote epithelial repair, and is upregulated in intestinal epithelial cells (IECs) from IBD patients. Moreover, loss of GSDMB was associated with decreased PDGFA expression, a growth factor known to induce FAK phosphorylation (11). These observations suggest the potential association between GSDMA/B and cell junction integrity in keratinocytes.

Here we study GSDMA and GSDMB in keratinocytes, taking into consideration the cell-type specific and unique features of keratinocytes regarding the unique and critical aspects of their cytoskeleton.

## Results

### Gasdermin A and Gasdermin B localize the cytoskeleton in keratinocyte

GSDMA is primarily expressed in stratum spinosum and granulosum layers of keratinocytes (9). While proliferating human keratinocytes express GSDMA protein, its mRNA levels increase upon differentiation. However, studies on the subcellular localization of GSDMA remain limited.

To investigate GSDMA localization in keratinocytes, we utilized immortalized human keratinocytes cell lines, N/TERT-1, and HaCaT cells which express all five gasdermins (GSDMA-E) (Fig 1-3), enabling direct comparison of their distinct activation pathways in a single cell type. Subcellular fractionation under basal conditions confirmed that GSDMD and GSDME are localized to the cytoplasm fraction in the N/TERT-1 and HaCaT cell line (Fig 1A, 1B), consistent with previous reports (12, 13). Notably, GSDMA and GSDMB were enriched in cytoskeleton fraction in N/TERT-1 cells (Fig 1A). In HaCaT cells, GSDMB was found to be predominantly expressed in the cytoplasm, whereas GSDMA exhibited a cytoskeletal localization similar to that observed in N/TERT-1 cells (Fig. 1B). This difference may be attributable to the relatively impaired barrier-forming capacity of HaCaT cells compared with N/TERT-1 and primary human foreskin keratinocytes (PHEKs) (14). Although both HaCaT and N/TERT-1 cells have been reported to exhibit abnormal epidermal stratification, aberrant expression of differentiation markers, and defective stratum corneum formation in 3D models, nevertheless, N/TERT-1 cells retain many of the key characteristics of primary human keratinocytes and have been shown to more closely recapitulate epidermal differentiation and stratified epithelial formation than HaCaT cells (15). Therefore, we believe that the results obtained using N/TERT-1 cells may more closely reflect the characteristics of human keratinocytes in the skin. We also examined Colo205 colon cancer cells, which also express both GSDMA and GSDMB. In Colo205 cells, GSDMA and GSDMB localized to the cytoplasm (Fig 1C), suggesting that cytoskeletal localization is dependent upon a specific aspect of keratinocyte cell biology.

**Figure 1.**
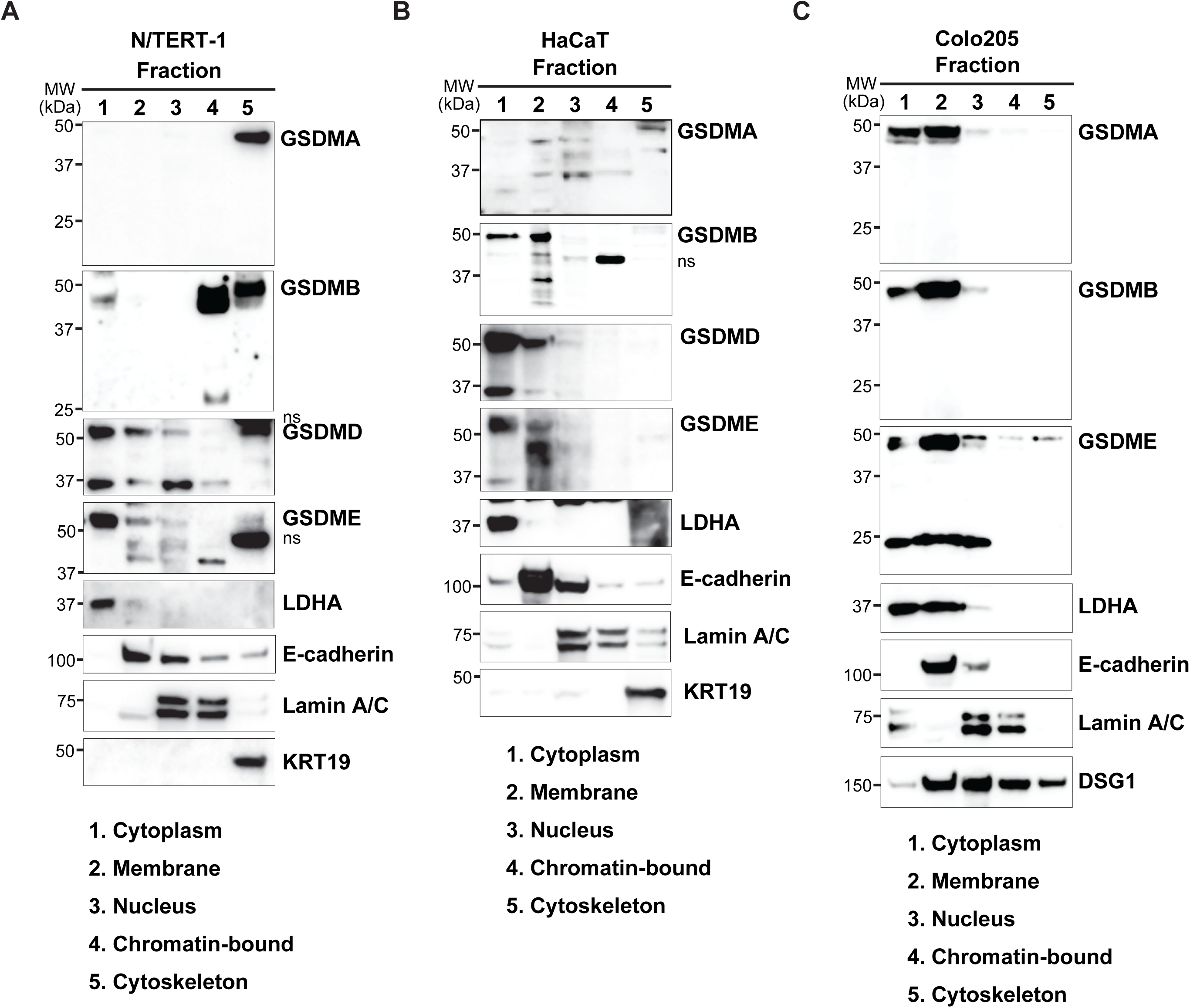
Gasdermin A and Gasdermin B localize the cytoskeleton in keratinocyte. (**A**) N/TERT-1, (**B**) HaCaT and (**C**) Colo205 cells were followed by subcellular fractionation into cytoplasmic, membrane, nuclear, chromatin bound, cytoskeleton as described in Methods. Each fraction was validated by immunoblotting with antibodies against LDHA, E-cadherin, Lamin A/C, DSG1, and KRT19. Numbers below the blots indicate individual purified fraction sample. Whole cell lysates were immunoblotted with the indicated antibodies. Molecular weights (MW, kDa) are represented on the left.

Together, these findings reveal that GSDMA and GSDMB specifically associate with the cytoskeleton in keratinocytes, distinguishing it from its cytoplasmic localization in other cell types and from GSDMD and GSDME.

### Gasdermin C localizes to the cell membrane and is not cleaved by caspase 8

Previous study has shown that the metabolite α-ketoglutarate (α-KG) promotes pyroptosis by inducing caspase 8-dependent cleavage of GSDMC in Hela cells (16). In response to α-KG, intracellular ROS levels increase, leading to oxidation of the membrane-localized death receptor DR6. A model was proposed where oxidized DR6 undergoes endocytosis and relocates to the cytoplasm, where it forms DR6 receptosome within which pro-caspase 8 is recruited and activated, and this active caspase 8 then cleaves GSDMC to trigger pyroptosis (16). Based on this model, GSDMC has been proposed to reside predominantly in the cytoplasm and to serve as a cytosolic substrate for caspase 8 (16).

To determine if this model is also observed in keratinocytes, we examined the subcellular distribution of endogenous GSDMC. Unexpectedly, in N/TERT-1 cells and HaCaT cells, GSDMC was detected mainly in membrane fraction, not the cytoplasm (Fig 2A, 2B). To assess whether membrane localization of GSDMC is restricted to keratinocytes, we performed the same analysis in the colon cancer Colo205 cell line and again found that GSDMC was predominantly present in the membrane fraction (Fig 2C). These data suggest that, contrary to the prevailing model, endogenous GSDMC is mainly associated with cellular membranes across multiple cell types.

**Figure 2.**
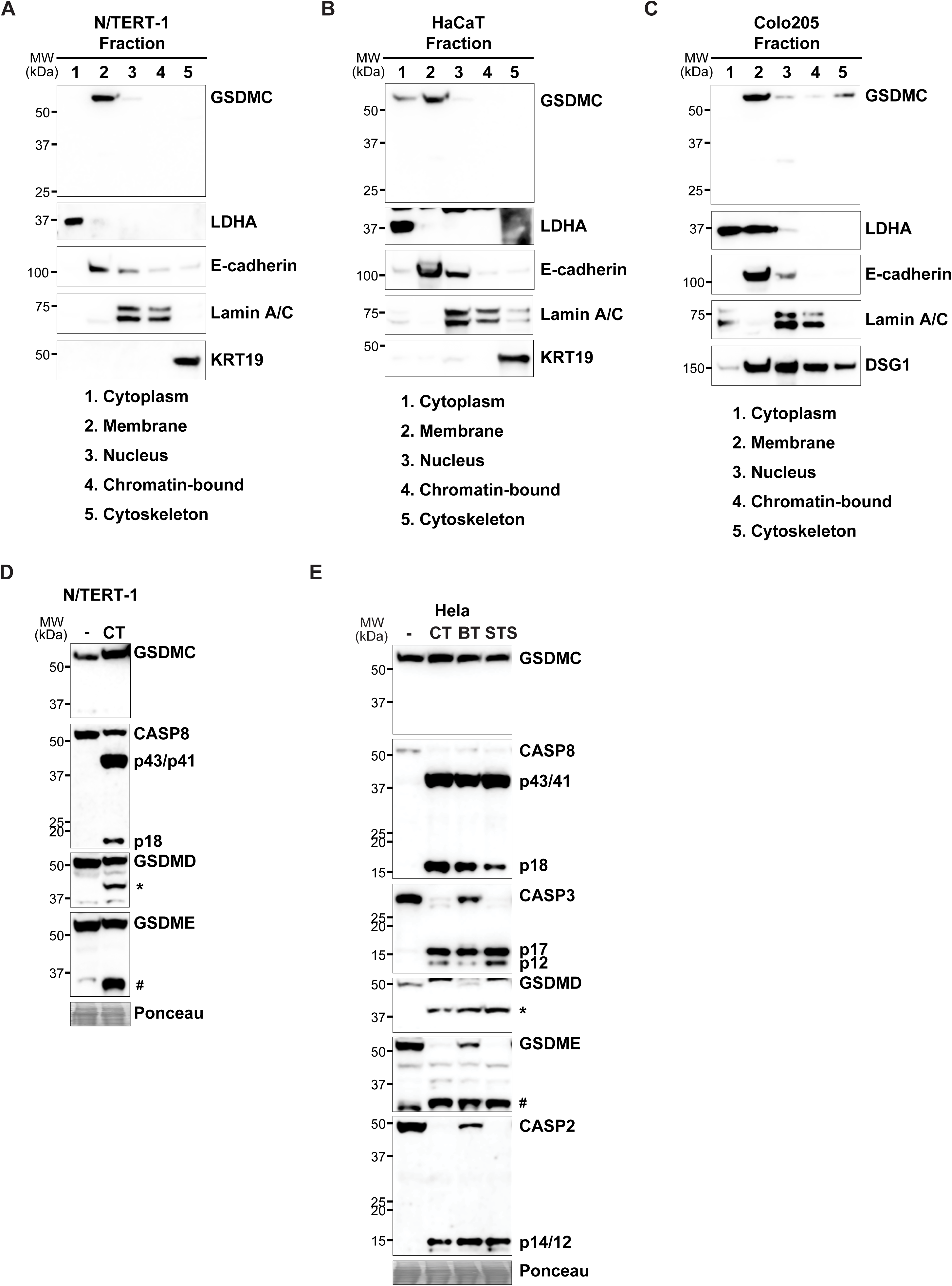
Gasdermin C localizes to the cell membrane and is not cleaved by caspase 8. (**A**) N/TERT-1, (**B**) HaCaT and (**C**) Colo205 cells were followed by subcellular fractionation into cytoplasmic, membrane, nuclear, chromatin bound, cytoskeleton as described in Method. Each fraction was validated by immunoblotting with antibodies against LDHA, E-cadherin, Lamin A/C, DSG1, and KRT19. Numbers below the blots indicate individual purified fraction sample. (**D**) N/TERT-1 cells were pretreated with cycloheximide (C, 100 µg/ml) for 30 minutes, followed by TNF (T, 100 ng/ml) treatment for another 24 hours. * indicates cleavage form of gasdermin D. # indicates cleavage form of gasdermin E. (**E**) Hela cells were pretreated with cycloheximide (C, 100 µg/ml), and BV6 (B, 10 µM) for 30 minutes, followed by TNF (T, 100 ng/ml), and staurosporine (STS, 10 µM) treatment for another 4 hours respectively. * indicates cleavage form of gasdermin D. # indicates cleavage form of gasdermin E. (**A-E**) Whole cell lysates were immunoblotted with the indicated antibodies. Molecular weights (MW, kDa) are represented on the left. (**D, E**) Equal protein loading was confirmed by Ponceau S staining.

Another study showed that PD-L1 translocated to the nucleus under hypoxia to enhance GSDMC transcription (17), and that TNF-induced caspase 8 cleavage of GSDMC shifts apoptosis to pyroptosis in breast cancer cells. To directly assess the ability of caspase 8 to cleave GSDMC, we treated N/TERT-1 cells with the well-established apoptosis inducer TNF plus cycloheximide (CHX). This treatment successfully activated caspase 8 as assessed by the presence of the p18 band (Fig 2D). However, GSDMC remained uncleaved despite caspase 8 activation (Fig 2D). As expected from prior reports, this activated caspase 8 cleaved and activated caspase 3, which cleaved the N-terminal domain of GSDMD (Fig 2D), which should prevent caspase 1 activation from driving GSDMD-dependent pyroptosis. Further, we also observed the known ability of caspase 3 to cleaved GSDME in its linker to promote pore formation, probably in settings where apoptosis fails (18) (Fig 2D). Observation of these GSDMD and GSDME fragments under these apoptosis conditions further confirms that our treatment successfully activated the caspase 8 to caspase 3 pathway. We further validated this observation in HeLa cells, a standard model for apoptosis studies, using multiple apoptosis inducers including IAP inhibitor, BV6 and staurosporine (STS) (Fig 2E). Although caspase 8 and caspase 3 were activated, alongside the expected cleavage of GSDMD and GSDME, GSDMC showed no evidence of cleavage.

Together, these data demonstrate that activated caspase 8 does not cleave GSDMC in keratinocytes and HeLa cells, indicating that we were unable to replicate earlier reports of caspase 8-dependent GSDMC cleavage (16).

### Granzyme H cleaves neither gasdermin A nor caspase 14

GSDMs are activated by proteolytic cleavage within the linker domain between N- and C-terminal regions. Inflammatory caspases (caspase 1/4/5/11) and executioner or apoptotic caspases (caspase 3/7/8) are well-established activators for GSDMs (19). Additionally, granzymes and other serine proteases such as neutrophil elastase have been shown cleave GSDMs to modulate pore-forming activity. Specifically, granzyme A expressed by cytotoxic T cells and NK cells can be delivered to target cells via perforin, where it cleaves GSDMB to trigger pyroptosis (20). In addition, granzyme B cleaves the linker region of GSDME to also promote pyroptosis (21), which might only occur after time in cells where apoptosis fails to complete. Among granzymes, granzyme H (GZMH) is highly expressed in human skin tissue (Human Protein Atlas) (Fig. S1) and human skin mast cells (22). We therefore hypothesized that GZMH might cleave the keratinocyte-specific GSDMA.

To test this hypothesis, we first generated active recombinant GZMH by incubating pro-GZMH with mouse capthepsin C (CTSC). Activation was confirmed by cleavage of DFF45, a well-established GZMH substrate (23) (Fig. S2A). As a positive control, we verified that SpeB from Group A *Streptococcus* cleaves GSDMA, consistent with previous reports (6, 24) (Fig. S2B). However, active GZMH failed to cleave GSDMA (Fig S3C). We also examined whether GZMH targets another keratinocyte-specific cell death protease, caspase 14 (25, 26), but observed no cleavage (Fig S4D).

Together, these data demonstrate that despite its expression in human skin, granzyme H does not cleave epidermal-specific proteins GSDMA or caspase 14. It remains possible that an unknown post-translational modification could enable these cleavage events.

### Gasdermin A is dispensable for vaccinia virus infection

Like most poxviruses, vaccinia virus (VACV) naturally infects keratinocytes in the skin and causes skin lesions. Because of this, VACV poses a significant risk to patients with atopic dermatitis (27) and serves as a well-established model for studying skin infection (28, 29). We therefore hypothesized that keratinocytes might detect VACV and activate GSDMA in response.

To test this hypothesis, we infected *Gsdma1/a2/a3* triple knockout mice (*Gsdma*^-^*^/^*^-^) (30) and wild-type littermates with VACV via dorsal flank scarification. Low-dose VACV infection induced erosive skin lesions that peaked at 3-5 days post-infection (dpi) in both genotypes (Fig S3A). *Gsdma*^-^*^/^*^-^ mice exhibited lesion progression patterns indistinguishable from wild-type controls. This phenotype persisted with high-dose VACV infections (Fig S3B). Neither genotype showed significant differences in survival rates through 15 dpi (Fig S3C), and skin lesion sizes were comparable at the peak (6 dpi) (Fig S3D).

These data demonstrate that GSDMA is dispensable for host control of VACV skin infections. This is in line with the ability of VACV to antagonize other regulated cell death pathways; VACV encodes multiple proteins that inhibit apoptosis, necroptosis, and pyroptosis (31).

### Calyculin A induces acantholytic cell death

Acantholysis is defined as the loss of cell-cell junctions in the skin. Loss of attachment to the ECM triggers a type of cell death called anoikis that is initiated by the loss of intracellular adhesion. We sought to determine the effects of acantholysis in keratinocytes in vitro, hypothesizing that disruption of the desmosome/hemidesmosome complexes would initiate a regulated cell death pathway. The PP2A inhibitors calyculin A and okadaic acid each can cause disruption of cell adhesions (32–35). Calyculin A treatment in N/TERT-1 cells caused cell death measured by Yoyo-1 dye uptake (Fig S4A-B). The activity of calyculin A to disrupt cell junctions can be observed as protein content of junctional components reduces, including those of the desmosome (desmogleins (DSG)), and adherens junctions (E-cadherin) (Fig S4C). Junction disruption has downstream effects that subsequently disassemble intermediate filament (IF) (vimentin) leading to reduced protein content (Fig S4C). Inhibition of PP2A causes excessive serine phosphorylation, resulting in disassembly of adhesion complexes. This disrupts FAK autophosphorylation and subsequent paxillin phosphorylation by the FAK–Src complex, leading to increased phosphorylation of integrin β4 (ITGβ4), which is indicative of adhesion complex disruption (36). Consistently, phosphorylation of ITGβ4 was increased but, phosphorylation of FAK and paxillin were decreased in late time points during calyculin A treatment (Fig S4D).

These data are in agreement with the known ability of calyculin A to cause cell junction disassembly and promote acantholysis.

### Calyculin A triggers translocation of GSDMA and GSDMB from cytoskeleton to plasma membrane

To determine whether desmosome/hemidesmosome disruption would change the subcellular localization of GSDMA and GSDMB, we treated N/TERT-1 cells with calyculin A to induce acantholysis. Subcellular fractionation revealed that GSDMA and GSDMB translocated from the cytoskeleton equally to both the cytoplasmic and membrane fractions upon calyculin A treatment (Fig 3A). To confirm whether this translocation is specific for GSDMA and GSDMB, we immunoblotted other GSDMs. GSDMC remained in the membrane as observed previously, while GSDMD and GSDME remained localized to cytoplasm. Notably, paxillin was dephosphorylated upon calyculin A treatment, consistent with acantholysis. To assess cell-type specificity, we examined Colo205 cells with calyculin A. Unlike keratinocytes, Colo205 cells showed no change in the localization of GSDMA and GSDMB, which were present in the cytosol and membrane fractions regardless of the treatment (Fig 3E). In contrast, GSDMC was localized specifically to the membrane fraction (Fig 3E).

**Figure 3.**
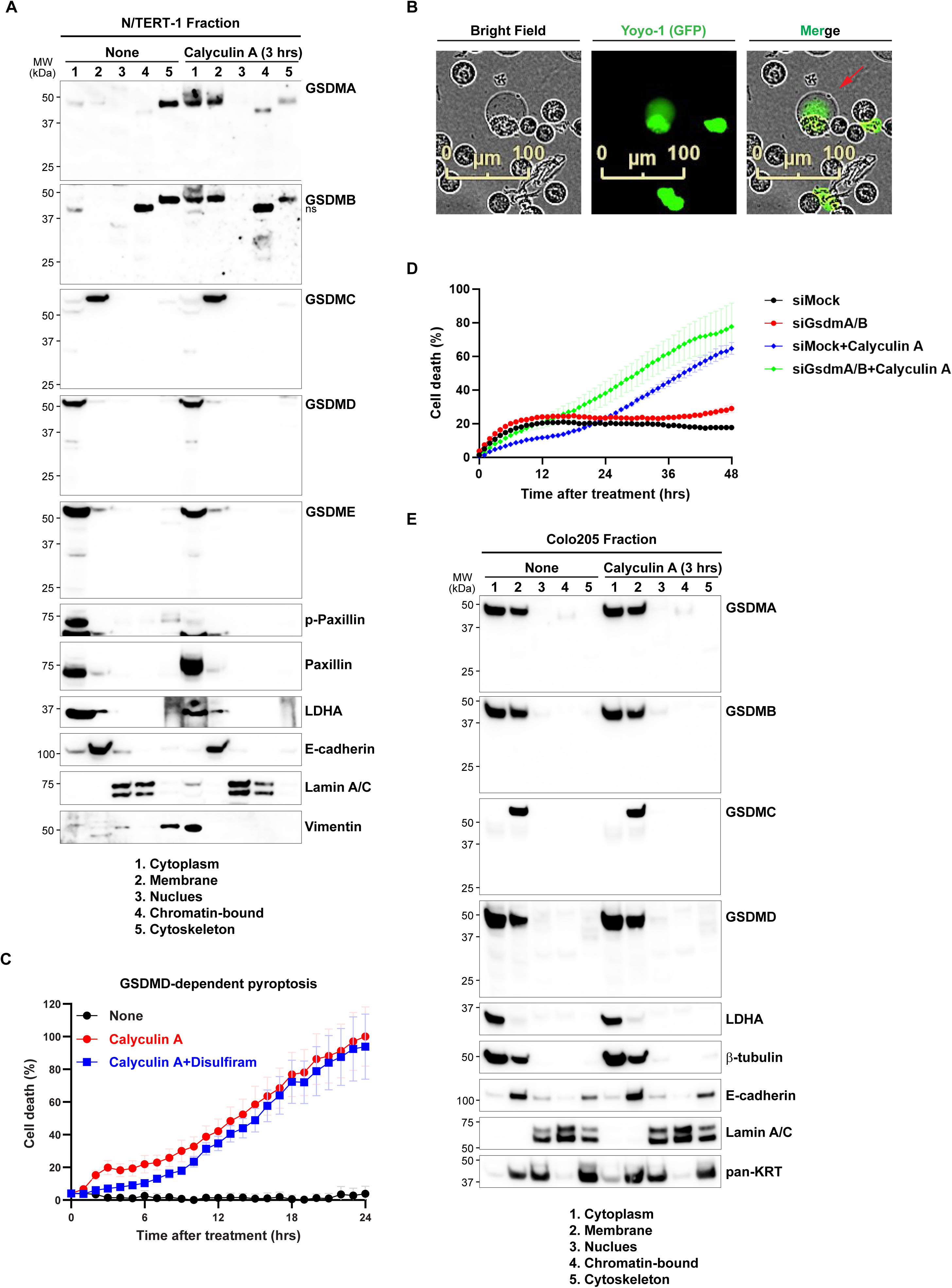
Calyculin A triggers translocation of GSDMA and GSDMB from cytoskeleton to plasma membrane. (A) N/TERT-1, (**E**) Colo205 cells were followed by subcellular fractionation into cytoplasmic, membrane, nuclear, chromatin bound, cytoskeleton as described in Method. Each fraction was validated by immunoblotting with antibodies against LDHA, β-tubulin, E-cadherin, Lamin A/C, Vimentin, and pan-KRT. Numbers below the blots indicate individual purified fraction sample. (**A, E**) Whole cell lysates were immunoblotted with the indicated antibodies. Molecular weights (MW, kDa) are represented on the left. (**B**) N/TERT-1 cells were treated with calyculin A (20nM) for 12 hours and stained with Yoyo-1 dye. Representative images from Incucyte live-cell imaging are shown. Red arrow indicates cell membrane ballooning. (**C**) N/TERT-1 cells were pretreated with Disulfiram (5 µM) for 30 minutes, followed by calyculin A (20 nM) treatment. (**D**) N/TERT-1 cells were transfected with either siMock or siGsdmaA/B and subsequently treated with calyculin A (20 nM). Cell death was quantified by Yoyo-1 dye uptake using Incucyte live-cell imaging. Graph represents percentage of cell death. Data are presented as mean ± SEM.

During calyculin A treatment, we did not observe cleavage of GSDMA or GSDMB. N-terminal fragments of GSDMs are known to translocate to the membrane and form a pore complex, which is hallmark of pyroptosis. A previous study demonstrated that full-length uncleaved GSDMD could localize to the membrane via palmitoylation without cleavage (37). This suggested the possibility that uncleaved GSDMA and/or GSDMB could open pores in the plasma membrane after calyculin A-induced membrane translocation. To determine whether calyculin A induced translocation of GSDMA and GSDMB triggers pyroptosis, we examined cell morphology by microscopy. Calyculin A treatment caused membrane ballooning and Yoyo-1 uptake, features that are characteristic of both acantholysis and pyroptosis (Fig 3B) However, no inhibition of cell death was observed following siRNA-mediated knockdown of GSDMA and GSDMB compared with the control (Fig 3D).

To further characterize the cell death pathway, we tested an inhibitor of GSDMD-dependent pyroptosis, disulfiram, which failed to suppress calyculin A-induced cell death (Fig 3C). The pan-caspase inhibitor, zVAD-fmk, also did not inhibit cell death (Fig S5A) and caspase 3/7 enzymatic activity was undetectable (Fig S5B). Similarly, other caspases were not activated as assessed by their cleavage (Fig S5C, S5D). Further, inhibitors of necroptosis (Nec-1) and ferroptosis (Fer-1) had no effect (Fig S5E, S5F). Taken together, these data confirm that calyculin A induces acantholysis, and calyculin A additionally drives translocation of GSDMA and GSDMB from the cytoskeleton to the membrane in keratinocytes.

### Endocytosis is required for calyculin A-induced acantholysis

Endocytosis is critical mechanism that internalizes disrupted cell junction proteins to facilitate recycling, which is a prerequisite to maintaining barrier function and tissue remodeling after junctional disruption (38, 39). Clathrin coats activate dynamin-mediated endocytosis, which is primarily responsible for the internalization of junctional proteins. We therefore hypothesized that clathrin-mediated endocytosis contributes to calyculin A-induced acanotholysis in keratinocytes.

To test this hypothesis, we pretreated N/TERT-1 cells with the clathrin inhibitor pitstop2 or the dynamin inhibitor dynasore for 30 minutes prior to calyculin A treatment. The characteristic membrane ballooning and cell death caused by calysulin A were markedly attenuated by pitstop2, resulting in morphology similar to untreated controls (Fig 4A). Dynasore produced comparable inhibition (Fig 4B). Pitstop2 suppressed calyculin A-induced cell death in a dose-dependent manners (Fig 4C). These data demonstrate that clathrin-mediated endocytosis is required for calyculin A-triggered acantholysis.

**Figure 4.**
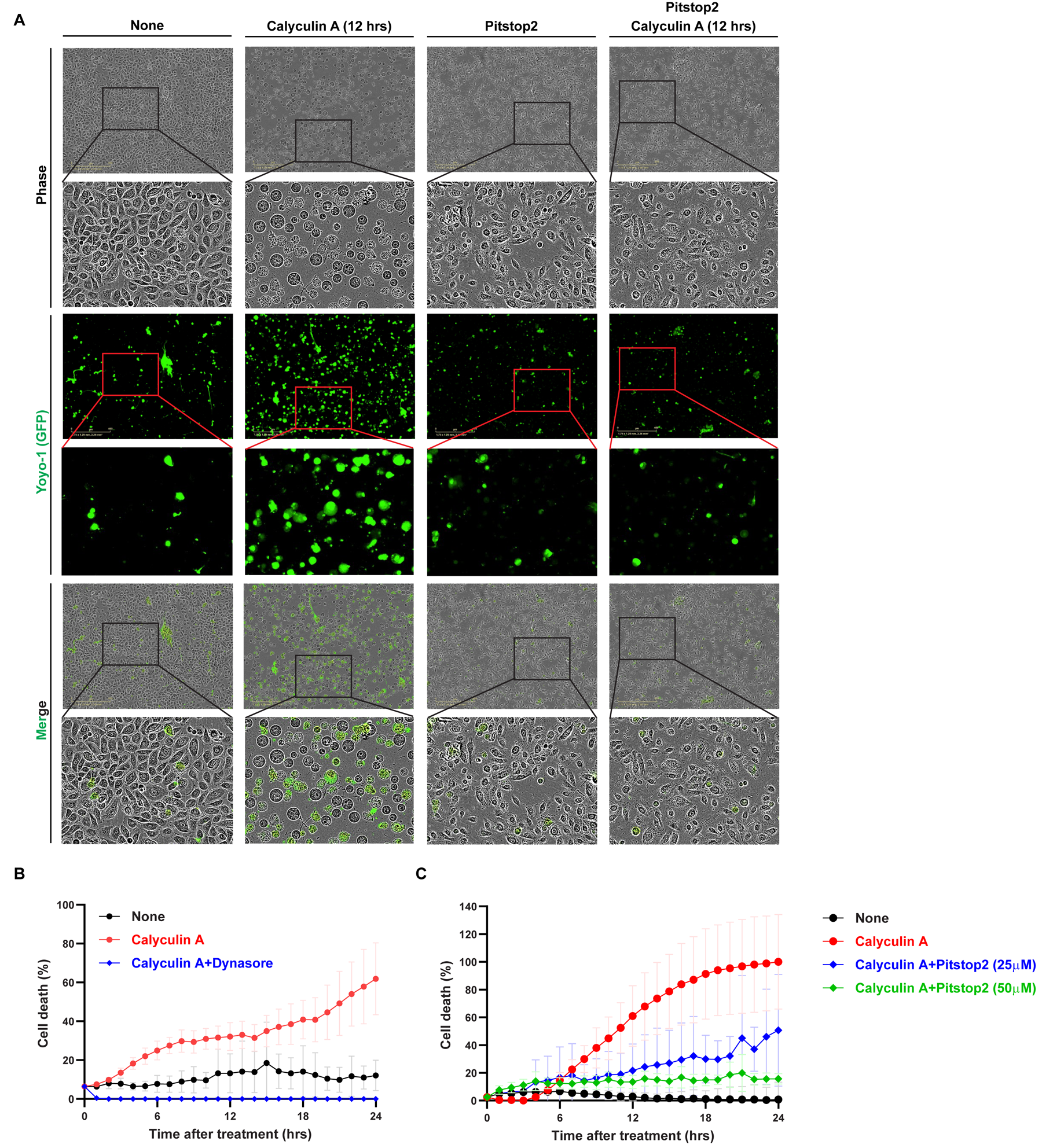
Endocytosis is required for gasdermin A and gasdermin B-dependent pyroptosis. (**A**) N/TERT-1 cells were pretreated with Pitstop2 (25 µM) for 30 minutes, followed by calyculin A (20 nM) treatment for 12 hours. Representative images from Incucyte live-cell imaging are shown after Yoyo-1 dye staining. (**B, C**) N/TERT-1 cells were pretreated with (**B**) Dynasore (100 µM) or (**C**) Pitstop2 at indicated concentration for 30 minutes, followed by calyculin A (20 nM) treatment. Cell death was quantified by Yoyo-1 dye uptake using Incucyte live-cell imaging. Graphs represent percentage of cell death. Data are presented as mean ± SEM.

### Calyculin A-induced acantholysis releases post-translationally modified keratin

Keratinocytes express several IL-1 family cytosolic cytokines, including IL-1β, IL-18, IL-33 and IL-36γ. Other cytosolic IL-1 family cytokines such as IL-33 and IL-36g do not require processing for their ability to signal through receptors, but they must be released from the cytosol in order to do so. Because calyculin A-induced acantholysis exhibited a membrane ballooning morphology similar to that observed during pyroptosis, we hypothesized that this process might also induce the release of inflammatory cytokines. To determine whether calyculin A-induced acantholysis could release IL-1 family inflammatory cytokines from N/TERT-1 cells, protein-containing media were concentrated by TCA precipitation and analyzed by immunoblotting. Unexpectedly, IL-1β, IL-33 and IL-36γ (40, 41) were not released during calyculin A-induced acantholysis (Fig 5A). Curiously, IL-18 was constitutively present in N/TERT-1 cells supernatants independent of calyculin A treatment, but it was not processed. Consistent with this, pretreatment with dynasore for 30 minutes prior to calculin A treatment had no effect on IL-18 release (Fig 5A).

**Figure 5.**
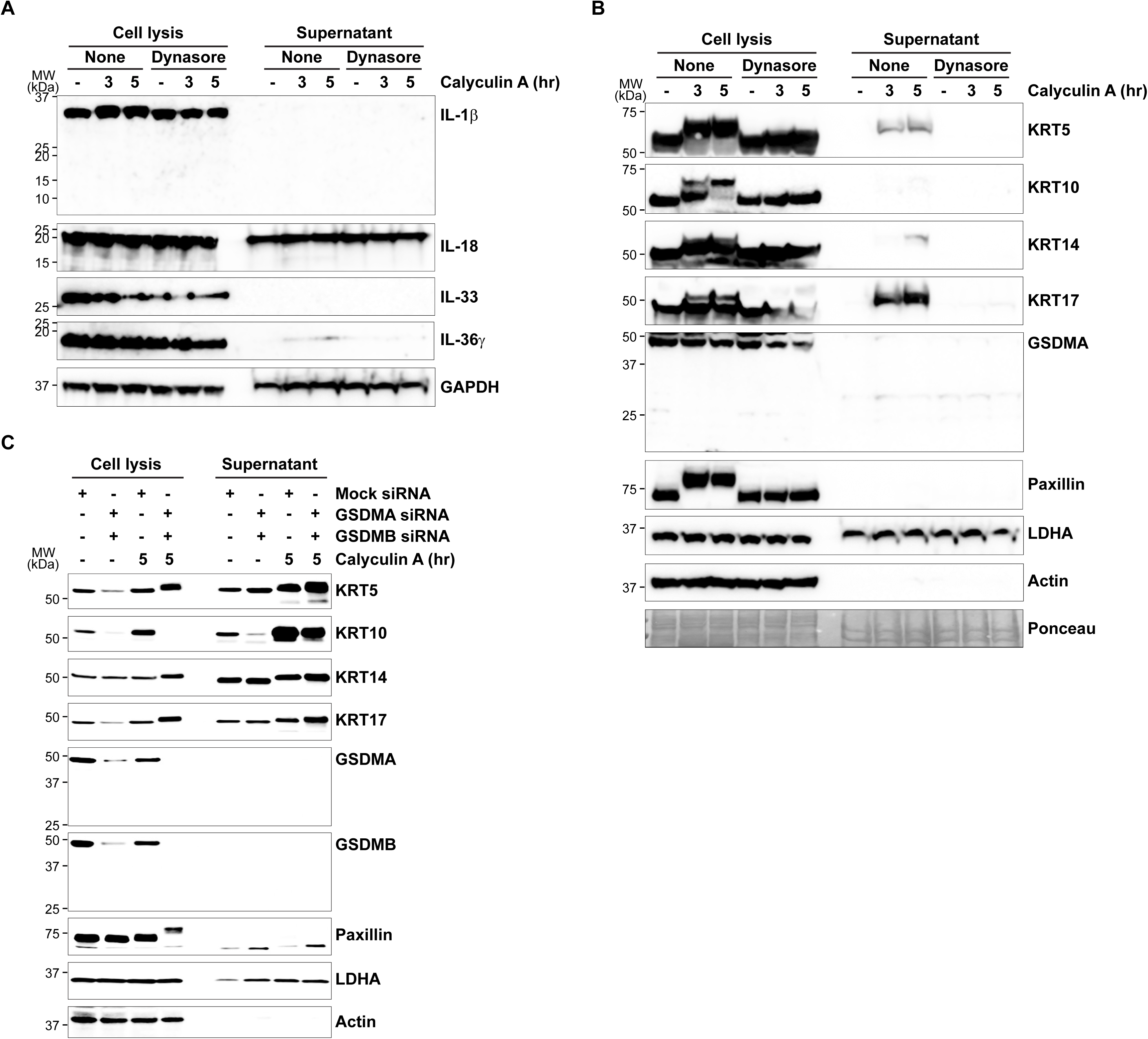
Calyculin A acantholysis releases post-translationally modified keratin. (**A, B**) N/TERT-1 cells were pretreated with Dynasore (100 µM) for 30 minutes, (**C**) N/TERT-1 cells were transfected with either siMock or siGsdmaA/B followed by calyculin A (20 nM) treatment for the indicated times. Cell lysates and supernatants separated as described in Methods. Whole cell lysates and supernatants were immunoblotted with the indicated antibodies. Molecular weights (MW, kDa) are represented on the left. Equal protein loading was confirmed by (**A**) GAPDH, (**B**) Ponceau S staining or (**C**) LDHA and Actin.

Previous studies have shown that lytic keratinocyte death and junctional or mechanical stress trigger keratin release (42, 43). We therefore hypothesized that calyculin A-induced acantholysis releases keratins. Consistent with this hypothesis, TCA-precipitated media from calyculin A-treated cells contained significantly elevated levels of keratins 5, 14, and 17 (Fig 5B). Additionally, keratin bands exhibited upward shifts, indicating post-translational modifications (PTMs) under calyculin A treatment. Previous studies have shown that keratins undergo PTMs upon mechanical or nonmechanical stress, associating with intermediate filaments (IFs) reorganization (44). To identify the keratin modification, we used inhibitors targeting specific PTMs. Keratins remained modified under 2-Bromopalmitate (2-BA) treatment, a palmitoylation inhibitor targeting S-palmitoyltransferase (Fig S6A). In addition, benzyl-α-GalNAc (O-glycosylation inhibitor), TAK-981 (SUMOylation inhibitor), and tunicamycin (N-linked glycosylation inhibitor) did not suppress keratin modification (Fig S6B-S6D). The keratin modification was independent on translational events (Fig S6E). The nature of this keratin modification remains to be elucidated. Finally, we used dynasore pretreatment, which eliminated keratin release (Fig. 5B). This confirms the requirement for endocytosis, suggesting a hypothesis that junctional complexes containing keratins are endocytosed, compartmentalized into multivesicular bodies and then exocytosed into the media.

To determine whether keratin release was associated with GSDMA and GSDMB, we performed siRNA-mediated knockdown of GSDMA and GSDMB. We found that keratin release induced by acantholysis was independent of GSDMA and GSDMB (Fig. 5C). These data demonstrate that calyculin A-induced acantholysis selectively released post-translationally modified keratin filaments.

## Discussion

GSDMA is cleaved by SpeB from Group A *Streptococcus* to form membrane pores that trigger pyroptosis (6, 24), while also playing a role in keratinocyte differentiation and cornification in models of skin barrier damage (45). GSDMB is cleaved by granzyme A from cytotoxic T cells or NK cells to induce pyroptosis via membrane pore formation (20). Here, we demonstrate that both GSDMA and GSDMB localize to the cytoskeleton in keratinocytes, which is a novel observation that provides structural evidence supporting their established associations with cell junction integrity.

Recent studies have reported that intraepithelial mast cells induce pyroptosis in intestinal epithelial cells (IECs) through the secretion of mMCP-1, which cleaves GSDMC (46). In addition, cathepsin S (CTSS) has been identified as another protease capable of GSDMC cleavage (47). In the present study, we did not attempt to validate mMCP-1 and CTSS, however, we were unable to validate the published ability of caspase-8 to cleave GSDMC (16). Interestingly, our data reveal the unexpected finding that full-length GSDMC is already localized to the plasma membrane in keratinocytes and other cell types. This observation raises the possibility that GSDMC may undergo constitutive palmitoylation prior to its activation by cleavage. Thus, unlike GSDMD and GSDME, which might require a two-step process involving activation by proteolytic cleavage as well as palmitoylation, GSDMC might require only a single step of proteolytic cleavage at the plasma membrane. While this hypothesis warrants further biochemical validation, our findings suggest different regulatory mechanisms governing GSDMD/GSDME compared to GSDMC-mediated pyroptosis.

Acantholysis refers to the loss of cell-cell adhesion or cell-matrix adhesion in keratinocytes due to disruption of intracellular junctions, resulting in rounded, detached cells that undergo anoikis (48). This pathological process is characteristic of autoimmune blistering disease such as pemphigus vulgaris (PV), where autoantibodies targeting desmogleins cause dismantling of desmosomes (49). Acantholysis can also be triggered by virus infections or mechanical stress. For instance, *Staphylococcus aureus* alpha-toxin binds ADAM10 to degrade tight junction proteins (ZO-1, ZO-2, occludin) and adherens junction components (E-cadherin), enhancing cytotoxicity (50, 51), while its protease SpeB cleaves GSDMA to induce pyroptosis (7). Our study demonstrates that calyculin A, a PP2A inhibitor, disrupts phosphorylation-dependent maintenance of junctional proteins, recapitulating acantholysis. Calyculin A-induced acantholysis, but this cell death was not prevented by knockdown of GSDMA and GSDMB. This result should be interpreted with a note of caution because it may be that the residual gasdermin proteins could be sufficient to form pores. Unfortunately, our efforts to knock out GSDMA in N/TERT-1 cells were not successful, but this approach warrants further investigation in future studies. Nevertheless, calyculin A did cause the translocation of uncleaved GSDMA- and GSDMB from the cytoskeleton to the plasma membrane, which could be part of a novel two step gasdermin activation mechanism. The calyculin A-induced acantholytic response requires clathrin-mediated endocytosis, as evidenced by suppression with dynasore and pitstop2. Clathrin-mediated endocytosis is essential for internalizing and recycling disrupted junctional proteins into endosomes during epithelial remodeling. These findings establish a junction disassembly-endocytosis-acantholysis axis, suggesting that the skin eliminate junctionally compromised keratinocytes through regulated acantholysis to preserve barrier integrity. This mechanism may extend to pathological contexts where mechanical stress or pathogen infection compromises junctions, requiring further investigation into the precise interplay between endocytic trafficking of junctional complexes and gasdermin translocation.

Recently, several studies have suggested alternative mechanisms to the well-established model in which gasdermins (GSDMs) are cleaved by proteases, allowing the N-terminal fragment to translocate to the plasma membrane and induce pyroptosis. Specifically, it has been reported that palmitoylation of GSDMD enhances its membrane targeting and thereby can promote pyroptosis even without cleavage (37, 52, 53). Interestingly, even a cleavage-deficient mutant (D275A) of GSDMD can undergo palmitoylation and translocate to the membrane in its full-length form, resulting in pyroptotic cell death, albeit less efficiently (37). In addition, under starvation conditions, activated ULK1 phosphorylates GSDMA at S353, and this phosphorylated full-length GSDMA can also move to the membrane and trigger pyroptosis without proteolytic cleavage (54). These findings indicate that GSDMs can induce pyroptosis not only in their cleaved forms but also as full-length proteins through specific post-translational modifications. Based on these observations, we propose that, in our experiments, GSDMA and GSDMB similarly translocated to the plasma membrane in response to cell junction disruption, possibly driven by unidentified post-translational modifications. Further studies are required to elucidate the nature of these modifications.

Mechanical or junctional stress disrupts the adhesion complex, which is subsequently internalized via endocytosis and packaged into recycling endosomes (39, 55). During this process, post-translational modifications (PTMs), such as phosphorylation, in keratin filaments, leading to disruption of the keratin network (43). PTM-modified keratins can be secreted extracellularly via the ER-Golgi pathway (56), while in apoptotic keratinocytes, caspase 6 cleaves keratin into fragments that are released into the extracellular space (42). These released keratins act as immunomodulatory DAMPs, promoting tissue repair signaling.

Additionally, keratin directly binds to DEC205/CD205 on dendritic cells (57), facilitating dead cell engulfment and antigen presentation (58). Our findings reveal that cell junction disruption triggers acantholysis in keratinocytes, and that this is different from classical GSDMD-mediated pyroptosis that is characterized by secretion of proinflammatory IL-1 family cytokines. Instead, this acantholysis model occurs concomitant with keratin release, which is could enhance dendritic cell-mediated immune responses.

Overall, our study proposes a novel mechanistic model that is distinct from the classical pyroptosis pathway. We demonstrate that calyculin A induces disruption of cell junctions, leading to acantholysis. During this process, GSDMA and GSDMB translocate to the plasma membrane in their full-length forms, without undergoing proteolytic cleavage. This finding supports the model in which keratinocytes undergoing junctional stress-driven extrusion are primed for a GSDMA- and GSDMB-dependent pyroptotic event, but a second stimulus may be required to trigger full pore formation. It is intriguing to speculate that full activation of a GSDMA or GSDMB pore could result in the specific release of an IL-1 family cytokine. Released keratins could subsequently binds to the DEC205/CD205 receptor on dendritic cells to promote dead cell engulfment and clearance. Through this work, we delineate a previously unrecognized route by which extruded keratinocytes arising from junctional stress are removed from the tissue.

Furthermore, we identify the PP2A inhibitor, calyculin A as a suitable tool to interrogate this pathway. These findings link the expression specificity of GSDMA in keratinocytes to a basic function that is unique to this cell type – the specific and specialized cytoskeleton that provides keratinocytes with structural integrity needed to form a barrier in the skin. Future studies will be required to define in greater detail how junctional stress signals drive GSDMA and GSDMB translocation to the membrane, and how endocytic processes counteract this translocation and modulate the ensuing pyroptotic response.

## Supporting information

Supplementary Figure 1. Granzyme H expression in organ

Supplementary Figure 2. Granzyme H does not cleave Gasdermin A and Caspase 14

Supplementary Figure 3. Gasdermin A is dispensable for vaccinia virus infection

Supplementary Figure 4. Calyculin A induces acantholysis by disrupting cell junction

Supplementary Figure 5. Identification cell death mode induced by Calyculin A

Supplementary Figure 6. Calyculin A modified keratin

## Acknowledgement

This work was supported by the NIH grants AI148302 and AI181815 (to E.A.M.).

---

**Supplementary Figure 1.** Granzyme H expression in organ. Organ-specific expression of granzyme H (GZMH) based on the data derived from Human Protein Atlas (proteinatlas.org)

**Supplementary Figure 2.** Granzyme H does not cleave gasdermin A and caspase 14. (**A-D**) Coomasie blue-stained gel shown in vitro cleavage assay using the indicated recombinant protein. (**A, C, D**) Recombinant protein human inactive granzyme H (GZMH) incubated with recombinant protein mouse active cathepsin C (CTSC) as described in Methods. GZMH and CTSC were serially diluted and incubated for 2 hours. Active form of GZMH was incubated with DFF45, (**C**) GSDMA, or (**D**) CASP14 for 30 minutes. (**A**) * indicates cleavage form of DFF45. Serial diluted concentration of SpeB incubated with GSDMA for 30 minutes. * indicates cleavage form of GSDMA.

**Supplementary Figure 3.** Gasdermin A is dispensable for vaccinia virus infection. (**A-D**) Wild type and gsdma^-/-^ mice were infected by vaccinia virus with indicated PFU. (**A, B**) Skin lesion sizes were quantified and plotted as graphs. Data are presented as mean ± SEM. (**C**) Graph represent percentage of survival. (**D**) Representative images showed skin lesions 6 days post infection. n indicates number of mice per group.

**Supplementary Figure 4.** Calyculin A induces acantholysis by disrupting cell junction. (**A-D**) N/TERT-1 cells were treated with calyculin A (20nM) for the indicated times. (**A**) Cell death was quantified by Yoyo-1 dye uptake using Incucyte live-cell imaging. Graph represents percentage of cell death. Data are presented as mean ± SEM. (**B**) Representative images from Incucyte live-cell imaging are shown after Yoyo-1 dye staining. (**C, D**) Whole cell lysates were immunoblotted with the indicated antibodies. Molecular weights (MW, kDa) are represented on the left. Cell junctions are categorized by the type on the right.

**Supplementary Figure 5.** Identification cell death mode induced by calyculin A. (**A, B, E, F**) N/TERT-1 cells were pretreated with (**A, B**) zVAD (20 µM), (**E**) necrostatin-1 (Nec-1, 50 µM), or (**F**) ferrostatin-1 (Fer-1, 10 µM) for 30 minutes, followed by calyculin A (20 nM) treatment. Cells were stained with Yoyo-1 dye, and cell death was quantified by Yoyo-1 dye uptake using Incucyte live-cell imaging. Graph represents percentage of cell death. Data are presented as mean ± SEM. (**C, D**) N/TERT-1 cells were treated with calyculin A (20 nM) for the indicated times. Whole cell lysates were immunoblotted with the indicated antibodies. Molecular weights (MW, kDa) are represented on the left.

**Supplementary Figure 6.** Calyculin A modified keratin. (**A-E**) N/TERT-1 cells were pretreated with (**A**) 2-Bromohexadecanoic acid (2-BA, 5 µM), (**B**) Benzyl-α-GalNAc (100 µM), (**C**) TAK-981 (10 µM), (**D**) Tunicamycin (10 µM), or (**E**) cycloheximide (CHX, 100 µg/ml) for 30 minutes, followed by calyculin A (20 nM) treatment for indicated times. Whole cell lysates were immunoblotted with the indicated antibodies. Molecular weights (MW, kDa) are represented on the left.

## Methods and Materials

### Antibodies and Reagents

GSDMA (ab230768), GSDMB (ab215729), CASP14 (ab174847), and p-ITGβ4 (ab29044) were purchased from Abcam, Calyculin A (101932-71-2), GSDMA (sc-376318), GSDMB (sc-101239), DSG1 (sc-137164), DSG2 (sc-365856), and IL-33 (sc-130625) were purchased from Santa Cruz, GSDMA (A22624), GSDMC (A21213), DSG1 (A9812), DSG2 (A19996, A21381), BP180 (A4808), CD151 (A1930), KRT5 (A11396), KRT10 (A4669), KRT14 (A19039), KRT17 (A23464), ITGβ1 (A23497), ITGβ4 (A4596), p-Paxillin (AP1156), Paxillin (A19100), IL-1β (A25874), IL-18 (A23076), IL-36γ (A23961), and GAPDH (AC002) were purchased from Abclonal, GSDMA (49307), GSDMD (97558), GSDME (84005), CASP1 (83383), CASP2 (2224), CASP8 (9746), CASP9 (9508), DSG3 (20483), pan-KRT (4545), Lamin A/C (2032), Vimentin (3390), p-Paxillin (69363), β-tubulin (86298), p-FAK (8556), FAK (62220), IL-1β (12242), and Actin (3700) were purchased from Cell Signaling, E-cadherin (610182), and Lamin A/C (612162) were purchased from BD bioscience, Mouse anti-rabbit IgG (111-035-144), and Goat anti-mouse IgG (115-035-062) were purchased from Jackson ImmunoResearch, Necrostatin-1 (4311-88-0), Necrosulfonamide (1360614-48-7), Ferrostatin-1 (347174-05-4), SP600125 (129-56-6), and Disulfiram (97-77-8) were purchased from Ambeed, Pitstop2 (HY-115604), staurosporine (HY-15141), zVAD (HY-16658B), and Zelenirstat (HY-147308), were purchased from MedChemExpress, Cycloheximide (66-81-9), and BV6 (533965) were purchased from Sigma-Aldrich.

### Cell lines

N/TERT-1 cells were kindly provided by Jenifer Zhang lab at Duke University. N/TERT-1 cells were cultured in keratinocyte media (17005042, ThermoFisher Scientific). HaCaT (300493, Cytion), and Colo205 (CCL-222, ATCC) cells were cultured in RPMI 1640 (11875093, ThermoFisher Scientific) including 10% fetal bovine serum, and Penicillin-Streptomycin (15140122, ThermoFisher Scientific) in 5% CO_2_ atmosphere at 37°C.

### Western blot

Cells were treated with conditions as indicated in the figure legends, harvested and lysed in RIPA buffer (10 mM Tris at pH 8.0, 1 mM EDTA, 0.5 mM EGTA, 1% Triton-X 100, 0.1% sodium deoxycholate, 0.1% SDS). Cell lysate were resolved by 5-20% gradient SDS-PAGE gels and transferred to nitrocellulose membrane (1620167, Bio-Rad). Membrane was blocked with 5% skim milk in TBS/T. Proteins were detected using ECL reagents (WBKLS0500, Sigma-Aldrich; or 34096, ThermoFisher Scientific), following the manufacturer’s instructions.

### Subcellular Fractionation

1-3 × 10^6^ of cells were plated onto 6 well plates or 6 cm dishes until 70-80% confluence. Subcellular protein fractionation kit (78840, ThermoFisher Scientific) was performed according to the manufacturer’s protocol. Fractions including cytoplasmic, membrane, nuclear, chromatin bound, cytoskeletal proteins were isolated. The purity of each fraction was validated by immunoblotting with antibodies as indicated in figure legends.

### SiRNA knockdown

N/TERT-1 cells were seeded in 6-well plates 1 × 10^6^ cells/well) and cultured overnight to reach 60–70% confluence. Transient knockdowns were performed using target-specific siRNAs (e.g., siGsdmaA/B) or non-targeting control siRNA (siMock) at a final concentration of 50 nM using Lipofectamine 3000 reagent (Invitrogen) in Opti-MEM Reduced Serum Medium, according to the manufacturer’s instructions. After 6 hours of incubation, the transfection medium was replaced with complete growth medium. Knockdown efficiency was validated 72 hours post-transfection via western blot analysis prior to subsequent chemical treatments.

### SiRNA sequence

The siRNA targeting human GSDMA:

Sense: 5’-rUrGrCrArArUrGrArUrArArCrArUrGrCrArArArCrCrUrUrC - 3’

Antisense: 5’-rGrArArGrGrUrUrUrGrCrArUrGrUrUrArUrCrArUrUrGrCrA - 3’ The siRNA targeting human GSDMB:

Sense: 5’-rGrArUrUrUrCrArUrArCrArUrGrGrArCrUrUrCrUrGrArUTA - 3’

Antisense: 5’-rUrArArUrCrArGrArArGrUrCrCrArUrGrUrArUrGrArArArUrCrCrA - 3’ SiMock

Sense: 5’ - rArCrGrUrUrUrUrUrUrCrUrGrrCrArCrArArUrArArUrArA - 3’

Antisense: 5’ - rUrUrUrUrArUrUrArUrGvGrUrGrArGrCrArGrArArArArArCrGrUrG - 3’

### Trichloroacetic acid (TCA) precipitation

To concentrate proteins from cell culture media, trichloroacetic acid (TCA) precipitation was performed. TCA was added to protein-containing media. Samples were incubated on ice for 30 minutes, followed by centrifugation at 10,000 x g, for 10 minutes at 4°C. The supernatant was aspirated. Protein pellets were resuspended in 2x SDS sample buffer. Proteins were resolved by SDS-PAGE and analyzed by immunoblotting.

### Mice

All mouse experiments were approved by the Duke Institutional Animal Care and Use Committee (IACUC). The UNC Animal Model Core generated Gsdma^-/-^ mice using CRIPSPR (30). Gsdma^-/-^ mice were born at the expected Mendelian ratios and exhibited no spontaneous skin abnormalities or dermatitis-like phenotypes prior to experiment.

### Mouse dorsal flank scarification

Wild-type and Gsdma^-/-^ mice (6-8 weeks old) were shaved on left dorsal flank. Hair gross and shaved skin were confirmed normal compared to wild-type mice. The following day, mice were infected with vaccinia virus on the shaved skin area under anesthesia conditions. The skin was gently scarified 30-40 times using a sterile 27G needle. Infected mice were maintained anesthetized until the viral suspension dried completely. Lesion sizes were measured daily for 14 days post-infection (dpi), and survival was monitored for 15 days.

### Vaccinia virus

The Western Reserve strain of vaccinia virus was amplified by passaging in BS-C1 cells for 72 hours. Cells were harvested following PBS washes twice, pelleted by centrifugation. Cell pellets were resuspended in PBS and subjected to freeze-thaw cycle 3 times. Virus titters were determined via plaque assay on Vero cells.

### Cell death assay and image

Cell death assay and image were monitored using the Incucyte S3 (Sartorius) live-cell imaging with 1µM of Yoyo-1 dye (Y3601, ThermoFisher Scientific) per wells. Green fluorescent, reflecting Yoyo-1 uptake by dead cells, quantified using the instrument’s integrated software.

### In vitro cleavage assay

Recombinant human DFF45 (NBP1-30202, Novus), human caspase-14 (MBS8120999, MyBioSource), SpeB (SPB-S5115-2000U, Acro Biosystems), human gasdermin A (RPE531Hu01, Cloud-Clone Corp), human granzyme H (GZMH, 1377-SE, R&D Systems), and mouse cathepsin C (CTSC, 2336-CY, R&D Systems) were used in the in vitro cleavage assays.Granzyme H (100 ug/ml) and cathepsin C (21 ug/ml) were incubated for 2 hours at 37°C to allow activation. Activated GZMH or SpeB was then incubate with indicated substrate as indicated in the figure legends for 2 hours at 37°C. Protein samples were resolved by 5-20% gradient SDS-PAGE gels and stained with Coomassie blue (1610435, Bio-Rad) according to the manufacturer’s instructions.

### Statistics

GraphPad Prism 8 software was used for statistical analysis. Error bars represent ±SEM. Unpaired t-tests or 2-way ANOVA were performed. p-values < 0.05 were taken statistically significant.

