## Supplementary figures and images for "Cell junction disruption drives translocation of gasdermin A and gasdermin B from cytoskeleton to plasma membrane during acantholysis"

### Supplementary Figure 1. Granzyme H expression in organ

**Supplementary Figure 1. Granzyme H expression in organ**

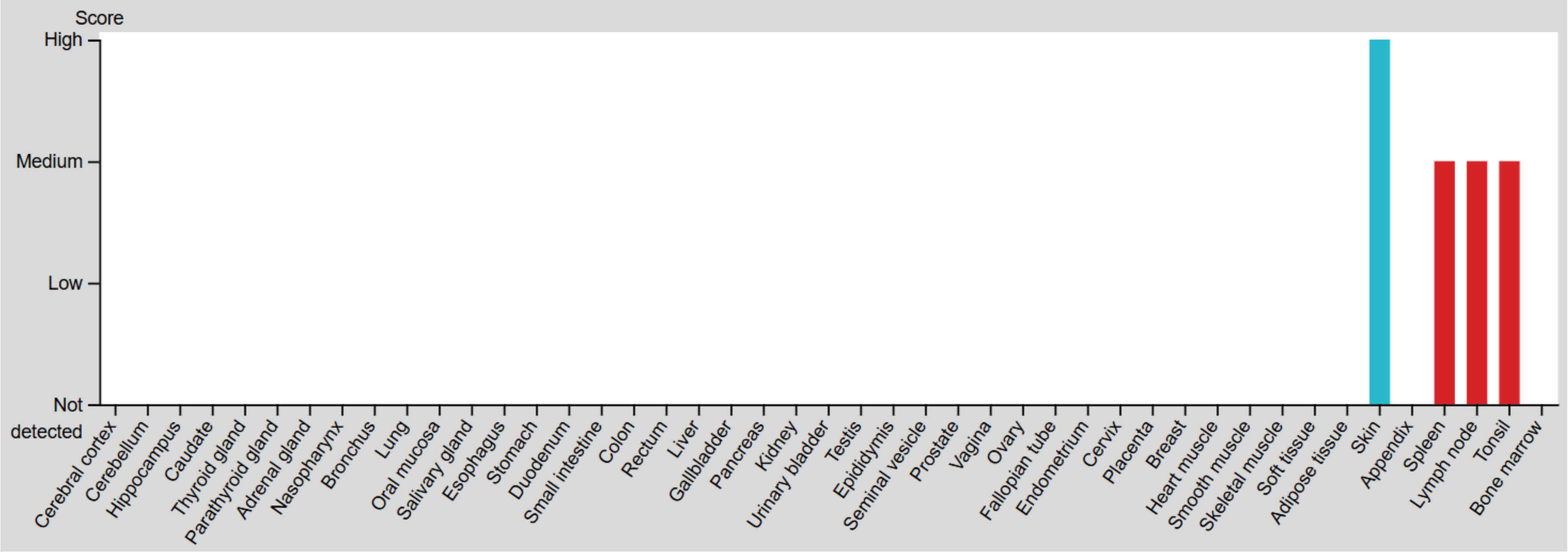

### Supplementary Figure 2. Granzyme H does not cleave Gasdermin A and Caspase 14

**Supplementary Figure 2. Granzyme H does not cleave gasdermin A and caspase 14**

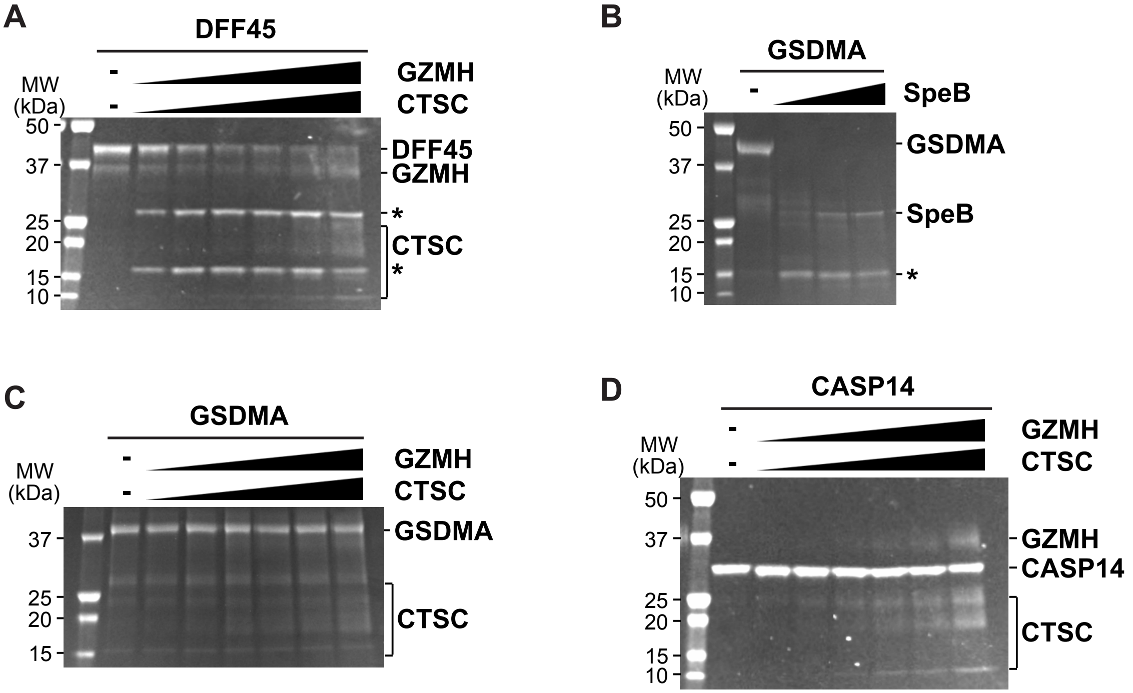

### Supplementary Figure 3. Gasdermin A is dispensable for vaccinia virus infection

**Supplementary Figure 3. Gasdermin A is dispensable for vaccinia virus infection**

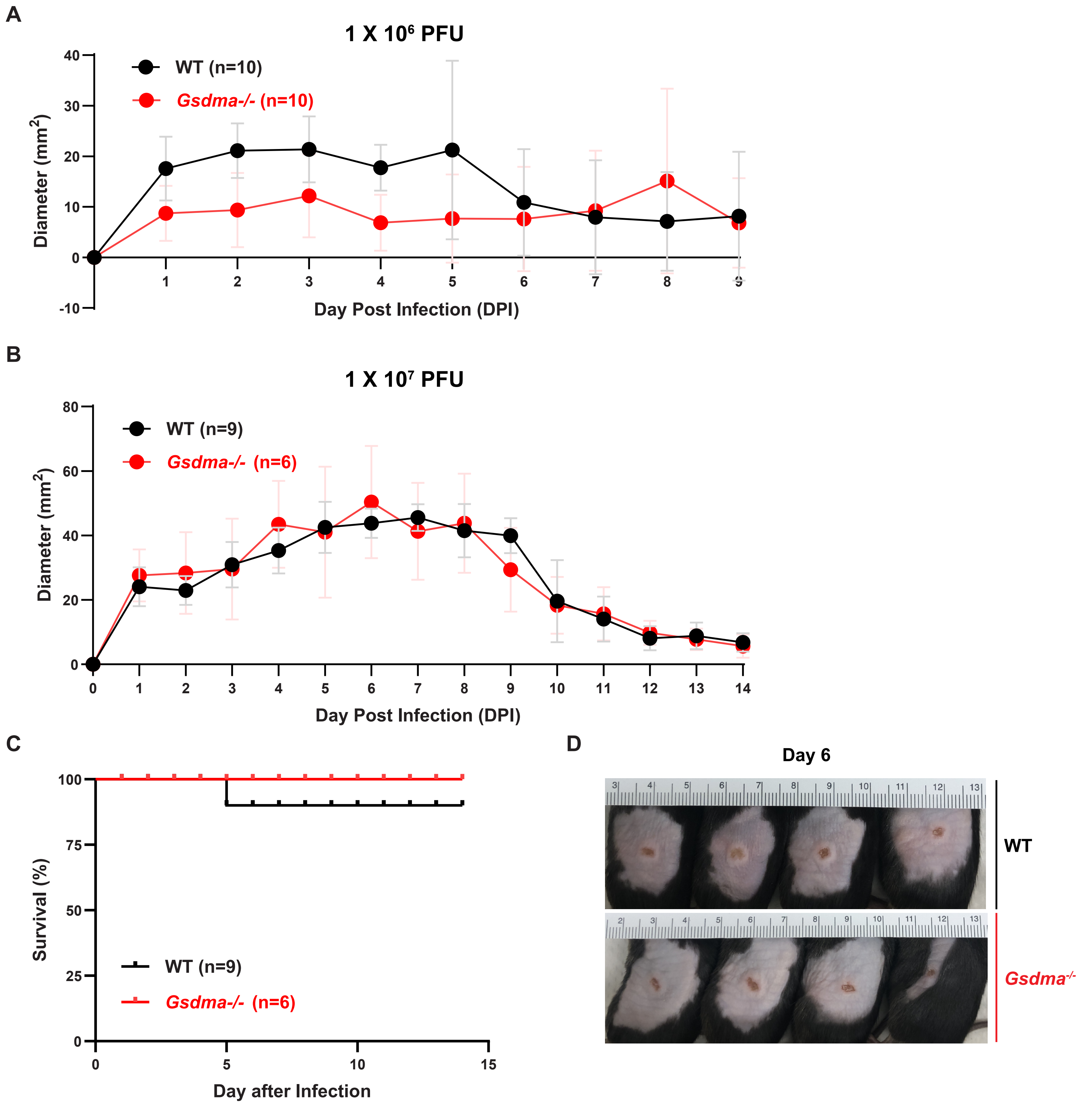

### Supplementary Figure 4. Calyculin A induces acantholysis by disrupting cell junction

#### Supplementary Figure 4. Calyculin A induces acantholysis by disrupting cell junctions

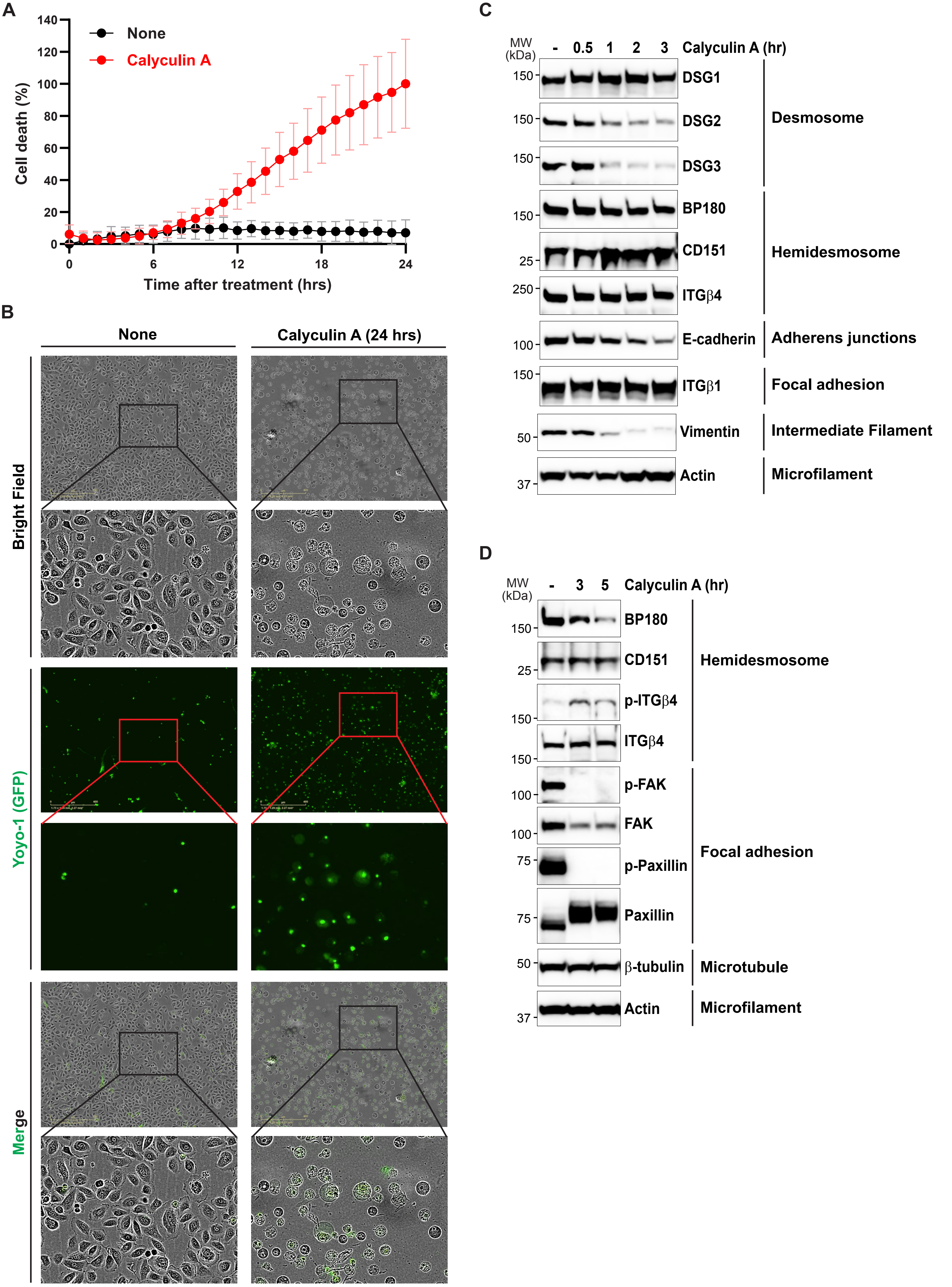

### Supplementary Figure 5. Identification cell death mode induced by Calyculin A

**Supplementary Figure 5. Identification cell death mode induced by calyculin A**

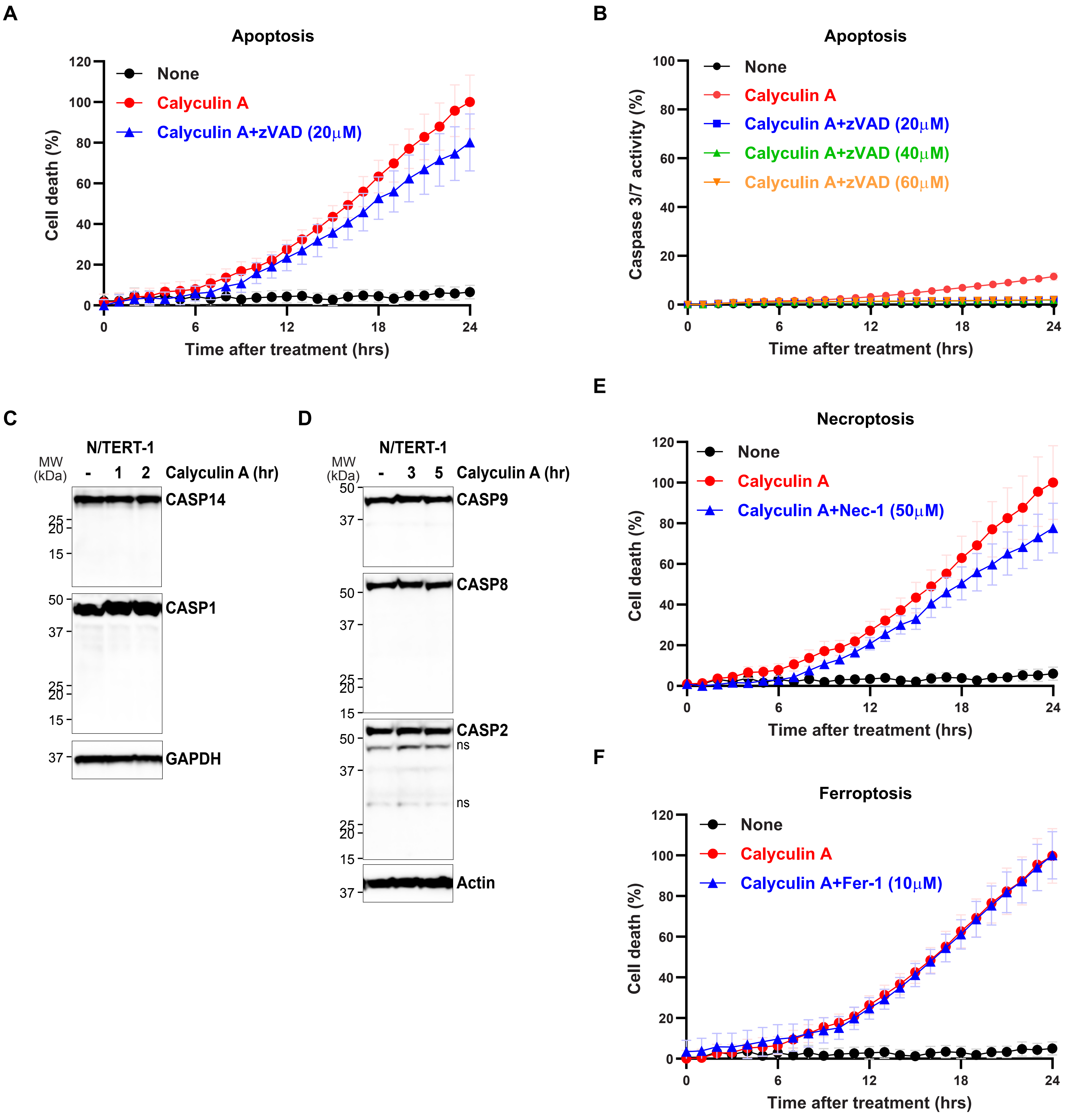

### Supplementary Figure 6. Calyculin A modified keratin

#### Supplementary Figure 6. Calyculin A modified keratin

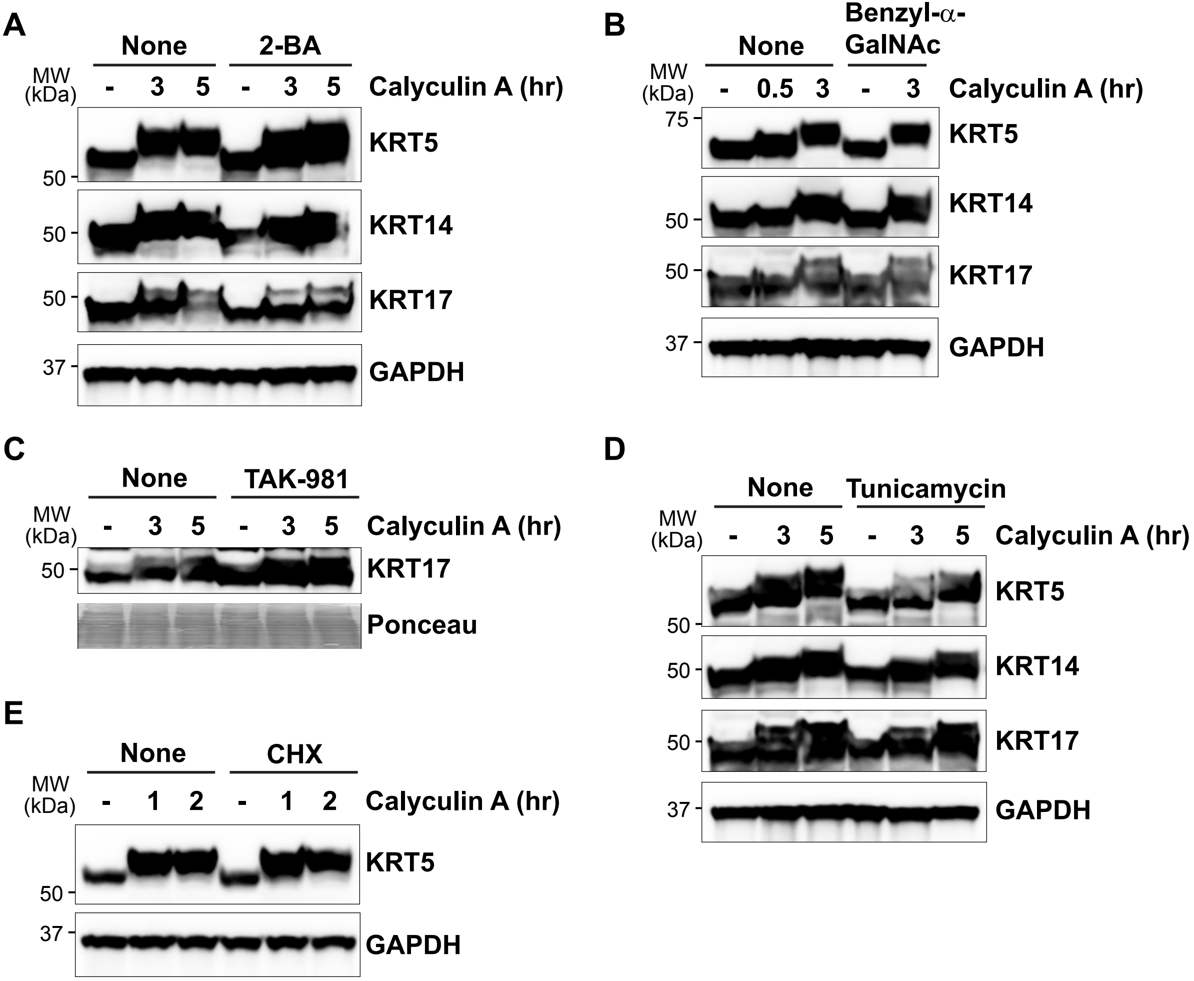
